# Partial overlap in the symptom profile induced by microglia activation and systemic inflammation

**DOI:** 10.64898/2026.03.02.709037

**Authors:** Priscila Batista Rosa, Silvia Castany, Andi Anderberg, Jack Tarakjian, Joost Wiskerke, Andreza Fabro de Bem, David Engblom

## Abstract

Microglial activation is a common feature of neurological and inflammatory diseases and may contribute to some associated symptoms. However, methodological limitations have made it challenging to identify the specific symptoms and behavioral consequences of selective microglial activation. In this study, we examined the spectrum of symptoms elicited by acute chemogenetic activation of microglia in mice and compared them to those induced by endotoxin-driven systemic inflammation. Both interventions upregulated inflammatory gene expression in the brain, reduced voluntary wheel running, and decreased self-care. Systemic inflammation additionally caused anorexia, weight loss, reduced motivation to work for palatable food, and impaired motor performance in the rotarod test—effects not observed with chemogenetic microglial activation. By showing that acute microglial activation reproduces certain motivational aspects of the sickness response while sparing other functions, the findings might shed new light on the contribution of microglia to symptoms and behavioral alterations during disease.

## Introduction

Microglia are the primary immune cells of the brain and play essential roles in both health and disease. In many neurological disorders, microglial function changes dramatically, leading to the release of inflammatory mediators. Such activation occurs in immunological brain diseases like multiple sclerosis, as well as in neurodegenerative conditions and stroke (1). While neurological diseases are typically characterized by symptoms arising from neuronal or circuit dysfunction, they are often accompanied by more generic disease symptoms such as loss of appetite, reduced motivation, and depression-like behaviors. Such symptoms also occur during systemic inflammation, where they are collectively referred to as the sickness response or sickness behavior (2).

Previous studies suggest that microglial activation can directly contribute to symptoms observed during immune challenges or in models of neurological and psychiatric disorders (3-7). However, it remains unclear whether such contribution is specific to only a few symptoms or of a broader relevance. Furthermore, many of the studies are based on deletion of microglial cells, which might lead to unspecific effects. Another approach to delineate the functional role of microglial cells is to activate them and investigate the molecular and behavioral outcomes. However, the specific symptoms and behavioral changes that microglial activation alone can induce have not been clearly defined. This knowledge gap largely reflects methodological challenges: inflammatory stimuli applied to the brain or general disease models typically activate multiple cell types, making it difficult to isolate the role of microglia. Fortunately, recent advances using designer receptors exclusively activated by designer drugs (DREADDs) have enabled selective microglial activation. These studies have shown that microglial activation in the spinal cord is sufficient to induce allodynia {Grace, 2018 #582; Saika, 2021 #671}, and activation in the striatum can produce aversion {Klawonn, 2021 #636}. However, a comprehensive assessment of the symptom profile following widespread microglial activation— and its relationship to the classic sickness response induced by lipopolysaccharide (LPS)—has not been conducted. Here, we used such tools to selectively activate microglia throughout the brain and describe the resulting sickness response. We also compared the profile of symptoms and behavioral changes to the well-established profile induced by systemic immune challenge with lipopolysaccharide (LPS). Our findings reveal that microglial activation recapitulates aspects of the sickness response to systemic inflammation but is more restricted – for example leaving appetite intact.

## Methods

### Animals

All experiments were conducted using both female and male mice; all animals were older than 6 weeks at the onset of the experiments, and the typical age at behavior assessments was 8-20 weeks. C57Bl6J mice (bred in-house or bought from Janvier Labs (Le Genest-Saint-Isle, France) were used for LPS injection experiments, and the transgenic line hM3Dq-Cx3cr1CreERT2 was generated from mouse lines from the Jackson Laboratories (*Cx3cr1creER;* stock #021160 and *CAG-LSL-hM3DGq;* stock #026220), bred in-house on a C57Bl6J background. Mice were kept under controlled conditions of humidity (40–60 %) and temperature (22 ± 1 °C) on a 12 h/12 h light/dark cycle (lights on at 7 A.M.) with food and water available ad libitum. The Animal Care and Use Committee at Linköping University approved all animal experiments.

### Drugs and injections

LPS (10 µg/kg; E. coli O111:B4, Sigma Aldrich; dissolved in sterile saline) or vehicle (saline) was administered intraperitoneally (i.p.). Clozapine-N-Oxide (CNO; Enzo Life Sciences) was given i.p. at 2 mg/kg (dissolved in sterile saline) for hM3Dq-activation. Cre-recombinase activity was induced by administration of tamoxifen (Sigma Aldrich) dissolved in a 10:1 sunflower seed oil/alcohol mixture. The mixture was injected i.p. at 2 mg for 5 days. To avoid the risk of DREADD expression in peripheral macrophages, we waited at least 4 weeks before administering CNO.

### Quantitative PCR

Mice were killed by asphyxiation with CO2 followed by cervical dislocation 2hours after injections of CNO or LPS. The striatum, hippocampus, and spleen were dissected, immersed in RNAlater (Qiagen; Hilden, Germany), and stored at −20 °C until analysis. RNA was extracted using a RNeasy Lipid Tissue Kit (Qiagen) following instructions from the manufacturer. A NanoDrop spectrophotometer (Thermo Fisher Scientific) was used to measure RNA concentration and quality. RNA samples included in the experiment showed A260/A280 and A260/A230 ratios > 1.8. The High-Capacity cDNA Reverse Transcription Kit (Applied Biosystems) was used for cDNA synthesis. Quantitative PCR was performed using TaqMan Gene Expression Master Mix (Applied Biosystems) together with the following TaqMan gene expression assays: Ptgs2;Mm00478374_m1, Il1b; Mm01336189_m1, Tnf; Mm00443258_m1, Il6; Mm00446190_m1, Cxcl10; Mm00445235_m1, and Ccl2; Mm00441242_m1. As an endogenous control, GAPDH; Mm99999915_g1 was used. Reactions were performed in a Real-Time 7900 Fast apparatus (Applied Biosystems). Relative quantification was carried out using the 2ΔΔCT method. Gene expression changes are reported as fold change relative to the respective control group.

### Immunohistochemistry

Mice were perfused transcardially with saline followed by buffered paraformaldehyde solution (4 %, pH 7.4). Brains were post-fixed for 24 h and incubated in 30 % sucrose-PBS solution until sinking. After freezing the brains, 30 µm coronal sections were cut on a cryostat (Leica Biosystems) and stored in anti-freeze solution at −20 °C until use. Immunolabeling for ionized calcium-binding adaptor molecule 1 (Iba1) was performed on free-floating sections. Briefly, the sections were incubated overnight with primary antibodies diluted in blocking solution: Iba1 (1:500; Fujifilm Wako), washed in PBS and incubated for 2h with appropriate secondary fluorescent antibodies with Alexa Fluor 488 (1:1000; all obtained from Invitrogen). Finally, the sections were mounted on glass slides using ProLong Glass Antifade Mounting agent (Thermo Fisher).

### Morphological analysis

Confocal images were acquired using a 40x objective Zeiss EC Plan-Neofluar. Iba1-positive cells in the striatum containing a single nucleus and no severed branches near the soma were selected. In all analysis, the experimenters were blind to the treatment. From each brain section, three to seven microglia were analyzed from each brain hemisphere in the striatal area, using ImageJ. Sholl analysis was used to quantify microglial morphology by measuring ramifications and cell body area. Concentric circles of increasing radius were applied around each microglial cell, and the number of intersections between the processes and the circles was counted to assess branching and territorial extension.

### Food intake and body weight measurements

Food intake was measured following brief fasting. On the day before testing, mice received half their regular daily food intake. On the experimental day, all remaining food was removed during the light phase, resulting in an 8-hour fast. Injections were done at the start of the dark phase. Food was returned 30 minutes after injections. Food consumption was measured 2.5 h after food was returned (3 h after injections) and again 12 h after injections. Body weight was recorded 12 h after injections.

### Sucrose self-administration

Three to four days prior to the start of the experiment, mice were food-restricted to half their regular daily intake until reaching approximately 90% of their initial body weight (sucrose pellets were provided in the home cage to habituate the animals to the reward). Mice were trained in operant conditioning chambers (Med Associates Inc.), where nose-poke responses resulted in the delivery of a sucrose pellet (Sucrose Dustless Precision Pellets, Bio-Serv ; Cat# F07595). After habituation to the chambers, animals underwent daily training sessions under a fixed ratio 1 (FR1) schedule until stable responding was achieved, correct responses were paired with a light cue signaling reward delivery. Subsequently, mice were trained under a fixed ratio 2 (FR2) schedule, in which two nose pokes were required to obtain one sucrose pellet. Once responding under FR2 became stable, animals were considered ready for experimental testing. Motivation for sucrose reward was evaluated using a progressive ratio schedule. During progressive-ratio (PR) sessions, the response requirement increased progressively according to the following formula: 5*e^(R*0.2)^-5, with R being number of rewards obtained + 1 (1,2, 4, 6, 9, 12, and so on). The highest ratio completed before the animal ceased responding for 30 consecutive minutes was defined as the breakpoint. The session was terminated either when this criterion was achieved or after a maximum duration of 1.5 h. Each breakpoint session was preceded by one or more FR2 sessions to re-establish stable performance before a new progressive ratio test was conducted.

### Voluntary wheel running

Voluntary wheel running was used to assess physical performance and motivation in effort tasks. Animals were single-housed and habituated in filtertop cages (37 × 21 × 18 cm) with free access to the vertical wireless running wheels (Med Associates; ENV-044V+DIG-804). The habituation period lasted 7–10 days, during which running activity progressively increased during the dark phase. After habituation, mice displayed stable and regular running activity across days, allowing experimental procedures such as injections to be performed. Wheel-running activity was continuously recorded. Injections were done one hour before the onset of the dark phase, as mice showed minimal activity during the light phase. The data from the injection day were compared to the data from the preceding day. The percentage of the activity was calculated using the mean of the 2 days before, with 100% as the reference. During the whole experiment, food intake and body weight were measured at 3–4-day intervals.

### Rotarod test

The rotarod test was used to investigate motor coordination and balance. The rotarod apparatus, with a rod 3.2 cm in diameter, was divided into 5 lanes, 8 cm each, fall height 30 cm (Med Associates, Inc.; ENV-574M). Before the start of the test, mice were allowed to habituate for 10 s at a constant speed of 4 rpm. Then, each mouse was tested for 5 min sessions, with a gradual acceleration rate from 4 to 40 rpm during each session. Latency to fall from the rotarod and end RPM were recorded for the same mice at 2, 4, 6, and 24 hours after injections.

### Open field test

The open field test was used to assess locomotor activity. In summary, the animals were placed in the center of an open-field arena (45 cm × 45 cm × 40 cm). Spontaneous locomotor activity and total time in the center were recorded and analyzed using EthoVision XT version 17 tracking software (Noldus) during a 10 min session.

### Sucrose splash test

The sucrose splash test was performed to evaluate self-care and anhedonia. Mice were placed individually in clear Plexiglas boxes (9×7×11 cm) and were sprayed on the back with a 10% sucrose solution, which naturally induces grooming. Animals were recorded for 5 minutes. The latency to initiate grooming and total duration of grooming behavior were recorded and quantified manually using EthoVision XT version 17 tracking software (Noldus).

### Statistical analysis

Statistical analysis was performed using GraphPad Prism version 10. Data are presented as mean ± standard error of the mean (SEM). Data normality was assessed using the Shapiro–Wilk test. Nonparametric tests were applied when the data were not normally distributed. Parametric tests were used when data were normally distributed or when the sample size was large.

Comparisons between two groups were conducted using the Mann–Whitney test or the t-test. For comparisons involving more than two groups, the Kruskal–Wallis test was used. When two categorical variables were analyzed, and data were normally distributed, a two-way ANOVA was performed, followed by Tukey’s post hoc multiple comparisons test. Statistical tests used, as well as sample sizes and other statistics, are stated in figure legends, the results section or in the supplemental statistical Table S1. A p-value < 0.05 was considered statistically significant.

## Results

To selectively activate microglia, we generated mice expressing an activating DREADD (hM3Dq) exclusively in microglia. The selectivity was achieved by combining the Cx3cr1-CreERT2 mouse line with a Cre-dependent DREADD-expressing line, an approach previously validated for microglial selectivity (3, 10). Based on our previous studies showing that microglia activation in the striatum contributes to inflammation-induced aversion, we activated the DREADDs with a low dose of CNO and quantified the levels of transcripts for Il1, Il6, Tnf, Ccl2, Cxcl10, and Ptgs2 in the striatum (Fig. 1A-G). Similar changes were seen in the hippocampus (Fig. S1A-C), indicating a widespread effect. The inflammatory genes studied were also upregulated following intraperitoneal injection of a low dose (10 µg/kg) of LPS (Fig. 1H-N), although the magnitude of induction varied between interventions. Morphological analysis revealed no statistically significant alterations in microglial structure in response to either intervention (Fig 1O-T), although a trend toward reduced ramification complexity was observed six hours after LPS injection. Collectively, these findings indicate that the DREADD-based approach elicited strong inflammatory signaling within microglia without triggering obvious morphological changes.

**Figure 1.**
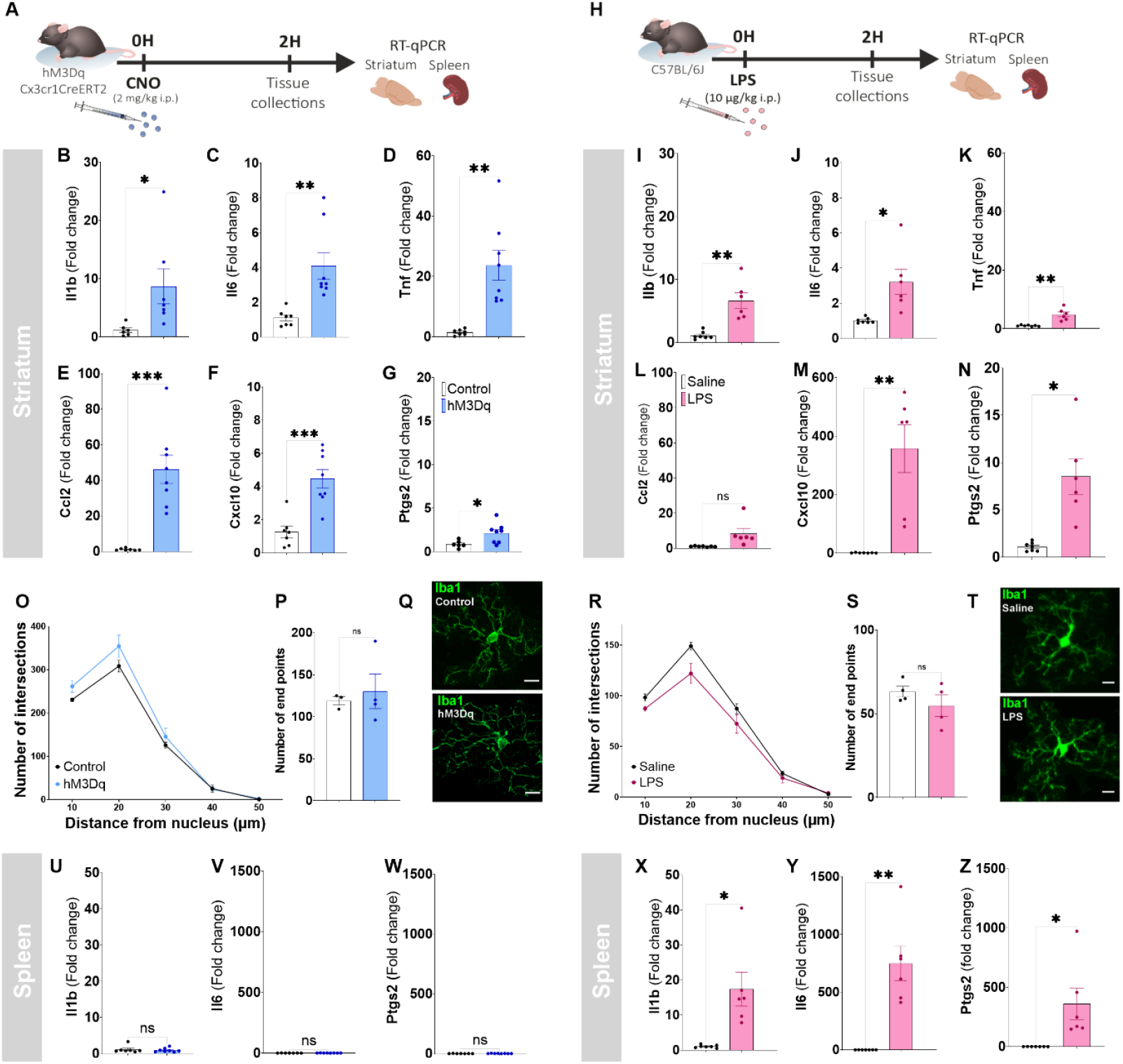
Characterization of DREADD-induced activation of microglia. The expression of inflammatory genes in the striatum was increased both by DREADD-induced activation of microglia (A-G; n = 7+ 8) and i.p. injection of lipopolysachharide (LPS; H-N; n = 7 + 6). No statistically significant changes in microglia morphology were seen after DREADD-activation (O-Q; n = 3 + 4) or LPS (R-T; n = 4 + 4), as can be seen in Sholl analysis of number of intersections (O, R) and end points (P, S). Representative images are shown in Q and T. DREADD-induced microglial activation elicited no induction of inflammatory genes in the spleen (U-W; n = 7 + 8), whereas i.p. injection pf LPS robustly did so (X-Z; n = 7 + 6). Statistical comparisons were done with t-tests except in O and R where two-way repeated measures ANOVA was used. Scale bar represents 10 µm. Data are shown as mean ± SEM. \**p* < 0.05; \*\**p* < 0.01; \*\*\**p*< 0.001.

To investigate if the DREADD-based microglial activation also led to systemic inflammatory signaling, we next investigated the expression of inflammatory genes in the spleen and liver. Notably, LPS caused a pronounced increase in inflammatory gene expression in the spleen, whereas DREADD-induced microglial activation did not induce any detectable changes (Fig. 1U-Z, Fig. S1D-F).

We next assessed whether DREADD-induced microglial activation elicited behavioral changes similar to those observed during systemic inflammation. Microglial activation did not reduce food intake 2.5 or 12 hours post-CNO injection (Fig. 2A-B), nor did it affect body weight (Fig. 2C). In contrast, LPS markedly decreased both food consumption and body weight (Fig. 2D, E) at the same time-points. Next, we investigated if microglial activation reduced the motivation to self-administer palatable food. Mice were trained to nose-poke for sugar pellets and after learning the task, their motivation was investigated using a progressive ratio protocol. LPS administered before the progressive ratio session nearly abolished self-administration of palatable food pellets, whereas no such effect was detected following microglial activation (Fig. 2F-J). Similarly, DREADD-induced activation of microglia did not affect the performance in the rotarod test (Fig. 2K, L), a test for coordination and balance, whereas LPS strongly reduced the performance in all sessions (Fig. 2M), indicating that microglial activation had no or limited effects on motor performance or learning in this paradigm.

**Figure 2.**
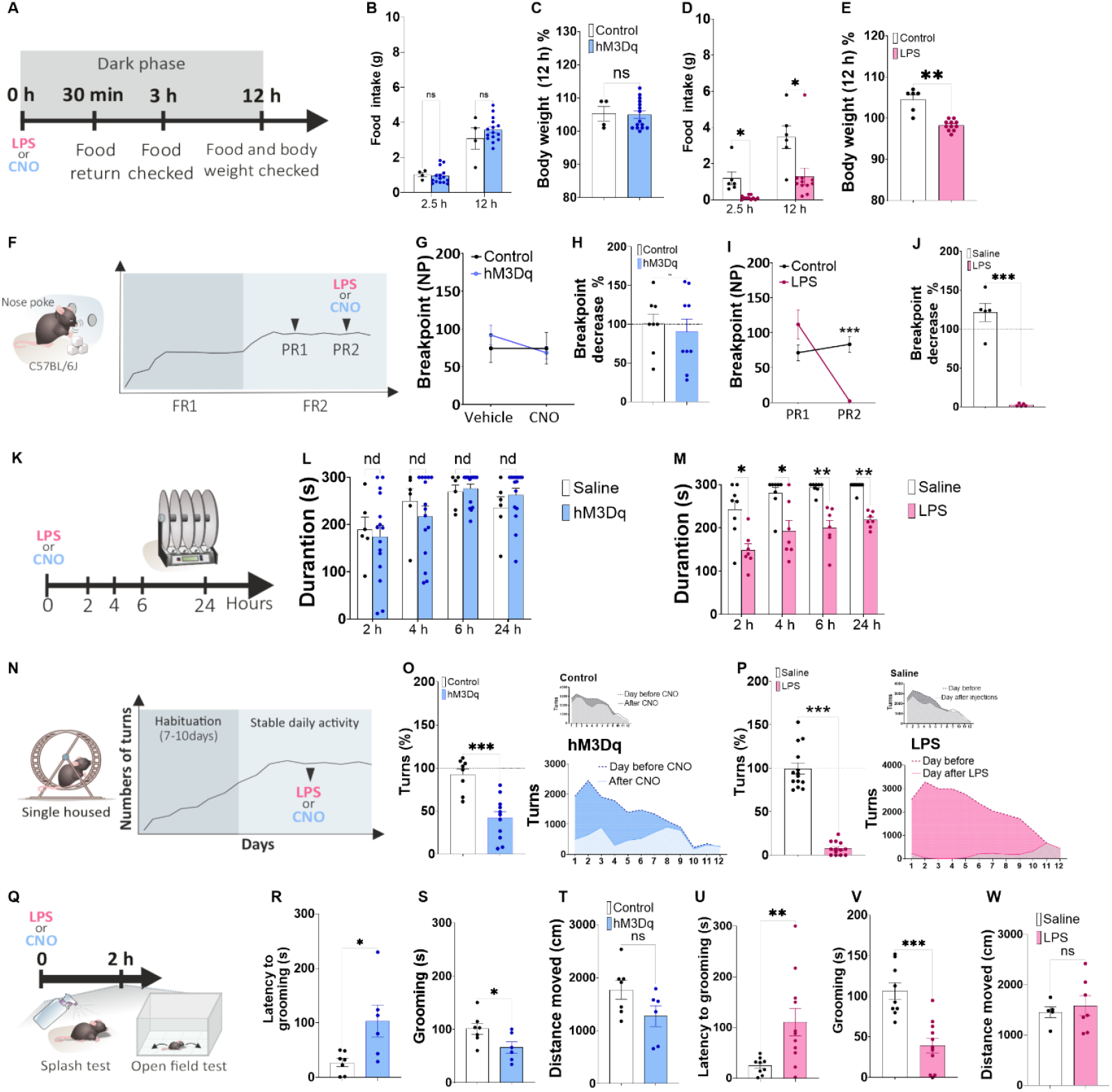
Behavioral consequences of microglial activation compared to systemic inflammation. Food intake and body weight (A) were not affected by DREADD-induced microglial activation (B, C; n = 4 + 15), but was robustly reduced by lipopolysaccharide (LPS) administration (D, E; n = 6 + 11). Self-administration of sugar pellets (F) was not reduced by DREADD-based microglial activation (G, H; n = 8 + 9), whereas it was almost abolished by LPS (I, J; n = 5 + 5). In the rotarod test (K), microglial activation did not induce any significant effect (L; n= 6 + 15), while LPS robustly reduced the level of performance (M; n = 7 + 7). Microglial activation reduced voluntary running in mice that had reached stable levels of running between days (N, O; n = 8 + 12). LPS also reduced voluntary wheel-running (P; n = 14 + 13). In the sucrose splash-test (Q), both DREADD-based microglial activation (P, Q; n = 7 + 6) and LPS (S, T; n = 9 + 11) increased latency to grooming (R, U) and reduced time spent grooming (T, V), without affecting locomotor activity in an open field-test (T; n = 7 + 6 and W; n = 5 + 7) performed right after the splash-test. Statistical analysis was done using two sample t-test (H, J, O, P, R - X), multiple t-tests (B, D), two-way ANOVA followed by Sidak’s multiple comparisons test (G, I), Multiple unpaired t tests (L), Multiple Mann-Whitney tests (M). Data are shown as mean ± SEM. \**p* < 0.05; \*\**p* < 0.01; \*\*\**p*< 0.001.

Next, we examined voluntary wheel running, a test sensitive to fatigue and reductions in motivation to move. After the animals had reached stable levels of performance between days, they were exposed to DREADD-induced microglial activation or LPS (Fig. 2N). Microglial activation led to a robust reduction in running (Fig. 2O), which was strongest in the first six hours, likely reflecting CNO pharmacokinetics. LPS also induced a strong and sustained decrease in wheel running (Fig. 2P). Finally, we investigated the effect of our interventions in the sucrose splash-test (Fig. 2Q), a test sensitive to apathy and reduced self-care. Microglial activation increased the latency to groom and decreased the time spent grooming, two hours post-CNO injection (Fig. 2R, S), without affecting spontaneous locomotion in an open field test conducted immediately afterwards (Fig 2T). Comparable results were seen after LPS administration (Fig. 2U-W). The findings demonstrate that isolated and acute microglial activation has clear behavioral consequences but does not replicate the full sickness syndrome (Fig 3).

**Figure 3.**
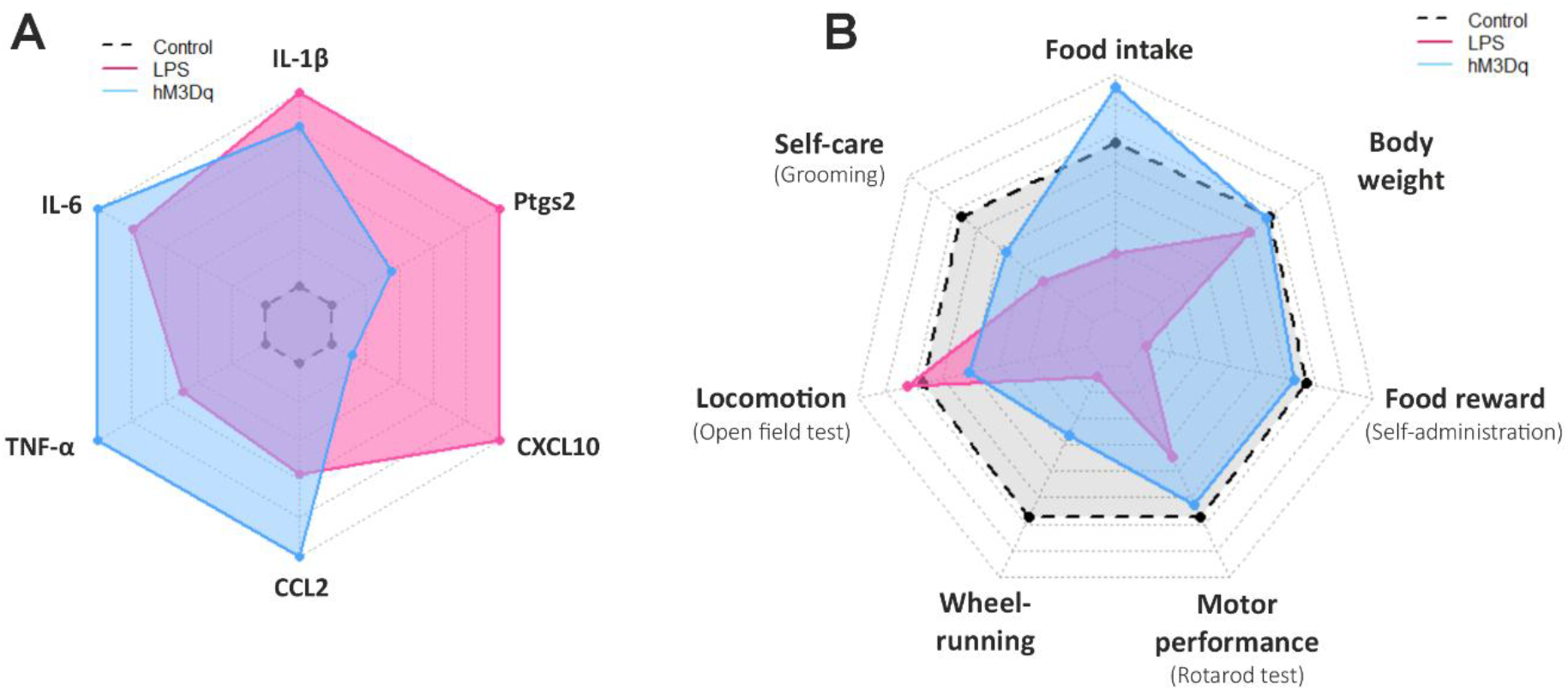
Comparison of cytokine-profiles and behavioral profiles after DREADD-induced microglial activation and systemic LPS. A: Levels of inflammatory mediator transcripts in the brain (striatum) after microglial activation (blue) and LPS (pink) on a logarithmic scale compared to saline injection (Control; grey). B: Behavioral profile after microglial activation or LPS. Levels are shown in percent of saline injected mice.

## Discussion

The present study demonstrates that selective chemogenetic activation of microglia produces a distinct behavioral and molecular profile that only partially overlaps with the classic sickness response induced by systemic LPS (see Fig. 3 for visualization). By utilizing the hM3Dq DREADD system, we were able to isolate the contribution of microglial signaling from the complex, multi-organ inflammatory response triggered by endotoxemia. More precisely, selective microglial activation was sufficient to reduce voluntary wheel running and impair self-care in the sucrose splash test, mirroring aspects of the sickness response. However, unlike systemic LPS, microglial activation did not induce anorexia, weight loss, or motor coordination deficits. This qualitative difference suggests that while microglia are central modulators of motivational states, potentially through interactions with dopaminergic or reward circuits, other symptoms like loss of appetite or reduced coordination may depend on broader systemic signals, such as peripheral cytokines acting on sensory afferents, circumventricular organs or the brain endothelium Consistent with previous studies (3, 8, 10), we observed a strong induction of inflammatory genes in the brain following CNO administration in mice expressing the Gq-coupled DREADD in microglia. Importantly, no corresponding induction was detected in the spleen, strongly suggesting that peripheral immune activation did not occur. This is noteworthy given a recent report indicating that activation of an inhibitory DREADD targeted to Cx3cr1-expressing cells may activate an unidentified peripheral immune cell population (11). Since the spleen is highly sensitive to peripheral immune signaling, it is unlikely that the pronounced behavioral effects observed in our DREADD mice were driven by systemic inflammation.

Furthermore, our molecular analysis revealed a “bias” in the inflammatory signature. LPS was significantly more potent in inducing endothelial-related mediators like *Ptgs2* (COX-2) and *Cxcl10* compared to the DREADD model. Given that the spleen remained unaffected in our DREADD-activated mice, this strongly suggests that the observed behavioral changes were driven by a strictly central microglial secretome. This highlights the utility of DREADDs as a precision tool to dissect the role of microglia in conditions where they are primary drivers of pathology, without the confounding effects of systemic multi-organ dysfunction. Since even a selective microglial activation inevitably influences other cell types over time, we restricted our analysis to single injections of CNO and early time-points. This could also explain the fact that we do not observe any obvious morphological changes in microglia after DREADD-activation. Previous studies (10, 13), as well as our own unpublished observations, indicate that repeated stimulation of microglia with DREADDs sometimes lead to unintuitive effects different from those induced by acute activation, complicating the interpretation of the results. Such effects can, for example, be caused by depletion of intracellular calcium stores in repeatedly activated microglia (13, 14).

The method used to induce microglial activation in this study is highly selective, at the cost of being quite artificial. This is difficult to avoid since we are not aware of any naturally occurring disorder in which an isolated microglial activation takes place. In this context, it is important to point out that microglia react differently dependent on the type of challenge, which means that the term microglial activation encompasses a number of different, but related, responses (15).

At the same time, most forms of acute microglial activations share common features like induction of cytokine genes, and our goal was to recapitulate such generic aspects. The decision to use Gq coupled DREADDs was based on literature suggesting that this is an efficient way to activate microglia (3, 8, 16) and that microglia express several Gq coupled receptors, including the P2Y6 receptor (17), that are important for their activation under physiological conditions.

Some of the mediators we found upregulated by DREADD-induced microglial activation have previously been shown to mediate microglia-dependent symptoms. Thus, allodynia elicited by activation of spinal microglia is inhibited by interventions with Il1 (8), and microglial Il6 is involved in inflammation-induced aversion (3). Such aversion is also dependent on prostanoids (19) and can be inhibited by selective deletion of microglial cyclooxygenase-1 (3). The model used here might be a good tool for studies on the mechanisms by which a generic form of microglial activation triggers associated symptoms, for example using specific gene deletions in microglia.

In conclusion, we show that microglial reactivity is a potent but selective driver of behavioral dysfunction. Acute microglial activation via a DREADD-based approach elicited a sickness response with a distinct profile compared to systemic inflammation. Microglial activation primarily affected motivated behaviors driven by non-food rewards whereas appetite-related measures and coordination were spared. These findings may inform future research on the contribution of microglia to symptoms and behavioral disturbances in inflammatory, neurological, and psychiatric disorders.

## Acknowledgements

This study was supported by the Swedish Research Council (2022-00952 and 2022-06568), the Swedish Brain Foundation (FO2022-0114, 2024-0263 and 2025-0299), the Knut and Alice Wallenberg foundation, Stiftelsen för Parkinsonforskning at Linköping University and Lions forskningsfond mot folksjukdomar. We thank the staff at the animal facility for technical assistance regarding animal breeding and care. We thank Maria Ntzouni and Vesa Loitto from histology and imaging facilities at Linköping Microscopy Unit for technical assistance. We also thank Daiane Engel and Sertan Arkan for their technical assistance and other members of the Engblom lab and Anders Blomqvist for discussions on design of the study and interpretation of the results, and input on the manuscript.

