## Supplementary material for "Partial overlap in the symptom profile induced by microglia activation and systemic inflammation": Fig S1

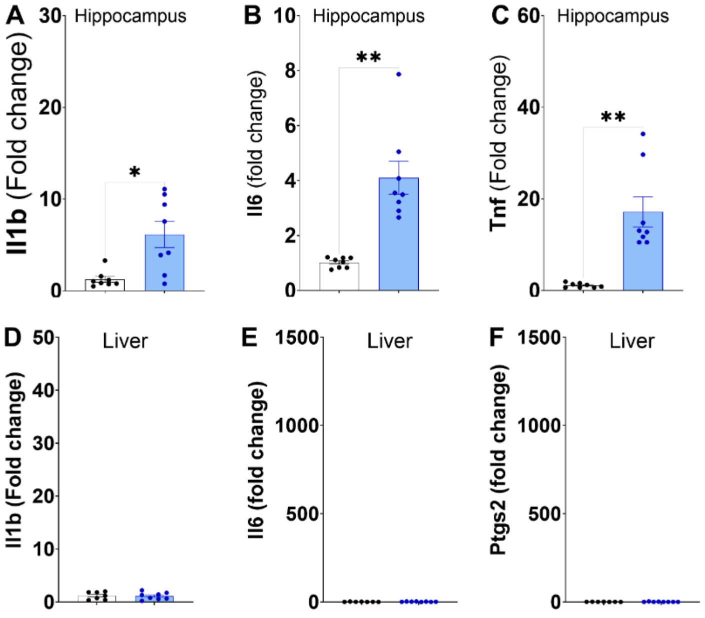


**Figure S1. Expression of inflammatory genes in the hippocampus and spleen in response to microglial activation.** The expression of inflammatory genes in the hippocampus was increased by DREADD-induced activation of microglia (A-C; n = 8 + 8), whereas no corresponding changes could be observed in the liver (D-F; n = 7 + 8).
