## Supplementary material for "Partial overlap in the symptom profile induced by microglia activation and systemic inflammation": Table S1

| Figure | Statistical test | Comparison | n | df | t/F | p | Figure display |
|---|---|---|---|---|---|---|---|
| 1B | Welch's t test | | 9 | 6,2180 | 2,4920 | 0,0456 | * |
| 1C | Welch's t test | | 15 | 7,7480 | 3,8100 | 0,0055 | ** |
| 1D | Welch's t test | | 15 | 7,0770 | 4,4670 | 0,0028 | ** |
| 1E | Welch's t test | | 15 | 7,0120 | 5,6720 | 0,0008 | *** |
| 1F | Welch's t test | | 15 | 4,8340 | 11,4600 | 0,0005 | *** |
| 1G | Welch's t test | | 15 | 9,2360 | 2,9110 | 0,0168 | * |
| 1I | Welch's t test | | 13 | 5,3710 | 4,4030 | 0,0059 | ** |
| 1J | Welch's t test | | 13 | 5,0660 | 3,0540 | 0,0278 | * |
| 1K | Welch's t test | | 13 | 5,1050 | 4,6380 | 0,0054 | ** |
| 1L | Welch's t test | | 13 | 5,0110 | 2,4490 | 0,0579 | ns |
| 1M | Welch's t test | | 13 | 5,0000 | 4,3490 | 0,0074 | ** |
| 1N | Welch's t test | | 13 | 5,0880 | 3,9070 | 0,0110 | * |
| 1O | 2way ANOVA | | 3+4 | | | | ns |
| | Radius x Groups | | | 4 | | 0,2088 | |
| | Radius | | | 4 | | <0,0001 | |
| | Groups | | | 1 | | 0,2656 | |
| | Subject | | | 5 | | 0,0019 | |
| 1P | Welch's t test | | 7 | 3,3520 | 0,5311 | 0,6286 | ns |
| 1R | 2way ANOVA | | 4+4 | 5,3180 | 2,7250 | 0,1800 | ns |
| | Radius x Groups | | | 4 | | 0,0638 | |
| | Radius | | | 4 | | <0,0001 | |
| | Groups | | | 1 | | 0,0818 | |
| | Subject | | | 6 | | 0,0022 | |
| 1S | Welch's t test | | 8 | 4,3730 | 1,1740 | 0,3005 | ns |
| 1U | Welch's t test | | 15 | 9,2110 | 0,4800 | 0,6424 | ns |
| 1V | Welch's t test | | 15 | 12,9300 | 0,7634 | 0,4589 | ns |
| 1W | Welch's t test | | 15 | 12,8400 | 1,0730 | 0,3029 | ns |
| 1X | Welch's t test | | 13 | 5,0110 | 3,3810 | 0,0196 | * |
| 1Y | Welch's t test | | 13 | 5,0000 | 5,0080 | 0,0041 | ** |
| 1Z | Welch's t test | | 13 | 5,0000 | 2,7210 | 0,0417 | * |
| 2B | Multiple unpaired t tests | 2.5 h | 4+15 | 17,0000 | 0,3585 | 0,7244 | ns |
| 2B | Multiple unpaired t tests | 12 h | 4+15 | 17,0000 | 1,1080 | 0,2831 | ns |
| 2C | Welch's t test | | 19 | 4,7490 | 0,1021 | 0,9228 | ns |
| 2D | Multiple unpaired t tests | 2.5 h | 6+11 | 5,0860 | 3,0250 | 0,0286 | * |
| 2D | Multiple unpaired t tests | 12 h | 6+11 | 10,3600 | 2,7930 | 0,0184 | * |
| 2E | Welch's t test | | 17 | 6,1410 | 5,1470 | 0,0020 | ** |
| 2G | 2way ANOVA, Šídák's multiple comparisons test | Vehicle | 8+9 | 30,0000 | 0,7977 | 0,6766 | ns |
| 2G | 2way ANOVA, Šídák's multiple comparisons test | CNO | 8+9 | 30,0000 | 0,2644 | 0,9573 | ns |
| 2H | Welch's t test | | 17 | 14,4300 | 0,5052 | 0,6210 | ns |
| 2I | 2way ANOVA, Šídák's multiple comparisons test | PR1 | 5+5 | 16,0000 | 2,1830 | 0,0866 | ns |
| 2I | 2way ANOVA, Šídák's multiple comparisons test | PR2 | 5+5 | 16,0000 | 4,3880 | 0,0009 | *** |
| 2J | Unpaired t tests | | 10 | 4,0450 | 10,1800 | 0,0005 | *** |
| 2L | Multiple Mann-Whitney tests | 2 h | 6+15 | 0,7000 | 42,0000 | 0,8361 | nd |
| 2L | Multiple Mann-Whitney tests | 4 h | 6+15 | 1,5170 | 38,5000 | 0,6308 | nd |
| 2L | Multiple Mann-Whitney tests | 6 h | 6+15 | -2,6830 | 33,5000 | 0,3529 | nd |
| 2L | Multiple Mann-Whitney tests | 24 h | 6+15 | -3,6170 | 29,5000 | 0,2305 | nd |
| 2M | Multiple Mann-Whitney tests | 2 h | 8+7 | 5,6250 | 7,0000 | 0,0137 | * |
| 2M | Multiple Mann-Whitney tests | 4 h | 8+7 | 5,4910 | 7,5000 | 0,0137 | * |
| 2M | Multiple Mann-Whitney tests | 6 h | 8+7 | 7,5000 | 0,0000 | 0,0003 | * |
| 2M | Multiple Mann-Whitney tests | 24 h | 8+7 | 7,5000 | 0,0000 | 0,0003 | * |
| 2O | Welch's t test | | 19 | 16,8200 | 4,9930 | 0,0001 | *** |
| 2P | Welch's t test | | 27 | 15,7100 | 13,4200 | <0,0001 | *** |
| 2R | Welch's t test | | 13 | 5,6660 | 2,4990 | 0,0489 | * |
| 2S | Welch's t test | | 13 | 10,8500 | 2,3150 | 0,0412 | * |
| 2T | Welch's t test | | 13 | 10,6000 | 1,8780 | 0,0882 | ns |
| 2U | Welch's t test | | 18 | 10,8100 | 3,1340 | 0,0097 | ** |
| 2V | Welch's t test | | 18 | 16,9900 | 4,8210 | 0,0002 | *** |
| 2X | Welch's t test | | 12 | 8,9270 | 0,5686 | 0,5836 | ns |
